# Hydrosphere occupancy modeling (ψ) and Akaike (AIC) habitat model selection of urban amphibian in West Java landscape

**DOI:** 10.1101/2021.04.23.441117

**Authors:** Adi Basukriadi, Erwin Nurdin, Andri Wibowo

## Abstract

Amphibians are animal that requires combinations of hydrosphere (riparian vegetation, water body) and vegetation (trees) microhabitats. In urban settings, those microhabitats are scarce and disappearing. One of areas that still have sufficient microhabitats to support amphibian populations is located in an 88.9 Ha urban forests of Universitas Indonesia Campus in West Java. Here, this paper aims to assess and model the several amphibian species occupancy (Ψ) with vegetation covers, riparian vegetation, and water bodies based on Akaike habitat selection (AIC) indices. The studied amphibian species include *Bufo melanosticus, Hylarana nicobariensis, Fejervarya limnocharis*, and *Polypedates leucomystax*. For modeling, 7 microhabitat models were developed and tested for amphibian species occupancy with covariates including vegetation cover, riparian vegetation, and water body microhabitats. The Principle Component Analysis (PCA) shows that the occupancies of amphibian were influenced mostly by the presences of water bodies followed by riparian vegetation, and vegetation covers. According to the values of Ψ and AIC, *Polypedates leucomystax* and *Fejervarya limnocharis* were species that have high occupancy in riparian vegetation microhabitats with Ψ values of −14.18 and −12.59. Likewise, *Hylarana nicobariensis* has an equal occupancy in vegetation, riparian vegetation, and water body microhabitats. While *Bufo melanosticus* shows high occupancy in vegetated microhabitats (Ψ = −14.18) rather than in riparian vegetation and water bodies (Ψ = −8.79). The combinations of riparian vegetation and water bodies show higher occupancy (Ψ = −8.00) rather than Ψ(vegetation cover+water body = −5.64) and Ψ(vegetation cover+ riparian vegetation = −4.36) combinations.

## INTRODUCTION

Habitat loss or degradation, climate change, and disease were known as attribute covariates that contribute to the rapid decline of amphibians worldwide. Amphibians are particularly vulnerable to those covariates present in hydrosphere habitats during both the aquatic and terrestrial life stages. Hydrosphere habitats including streams and ponds are suitable microhabitats for eggs, larvae, and tadpoles for months or even a full year. Juvenile amphibians are often dispersed from their natal ponds during metamorphosis. Semlitsch & Bodie (2003) and Becker et al. (2007) stated the quality of the environment with its attribute covariates both in and around breeding sites, is likely to be an important determinant of amphibian persistence in greatly altered urban landscapes.

In urban landscapes, urbanization and progressive changes in land use are considered to exert some of the strongest influences on amphibian populations worldwide. Adverse human activities threatening amphibian species including urban expansion, development of dense road networks without provision of compensatory solutions, and increased runoff contribute to the widespread loss or degradation of habitats, water pollution, isolation, and other unrecognized threats. Those conditions may lead to the drastic decline or even extinction of amphibian local populations of the most vulnerable species. Collins & Storfer (2003) and Pounds et al. (2006) noticed that given the spread of invasive species and pathogens including chytrid fungi and ranaviruses, and combined with climate change, the situation of amphibian populations appears to be critical worldwide. The observed decline of this group of vertebrates may constitute a symbol of hydrosphere biodiversity loss due to anthropogenic pressure.

This condition require novel approach to assess how those covariates influence the amphibian populations. One of approach is using the occupancy modeling. This modeling method has emerged as a tool to assess presence that is well-suited to large landscapes or patchy habitats. Occupancy models allow variable occupancy (or likelihood that a patch is occupied by a species of interest) and determinant covariates to be calculated for species within a habitat patch. Occupancy data is represented as a binomial representation of presence or absence based on a minimum of two repeat visits within an ecologically defined period of time. A distinguish feature of occupancy model is the ability to incorporate variation in species occupancy that may result from survey specific or site specific covariates that can also affect occupancy. Site specific covariates are potential for modeling the amphibian occupancy.

Current data on the occupancy of amphibians in hydrosphere of urban landscape mainly in South East Asia are still insufficient. Knowledge about amphibian populations living in SE Asia biggest cities appears important to conserve amphibian populations. In this paper, this study presents a complex study of amphibian occupancy in the 88.9 Ha urban forests of Universitas Indonesia Campus in West Java. The method to determine the amphibian occupancy was using occupancy modeling (Ψ) and Akaike (AIC) habitat model selection methods.

## MATERIALS AND METHODS

### Study area

This study was conducted in an 88.9 Ha urban forest of Universitas Indonesia Campus in West Java in longitude of 106.81-106.83 East and latitude of 6.34-6.37 South (Figure 1). The urban forest has high NDVI levels closed to 1 indicating dense vegetation covers. The moisture levels in urban forest also have values equal to 1 indicating high water contents in these areas. The hydrosphere in urban forest including a lake located in the central of the forest. Vegetation in the banks of the lake has contributed to the riparian vegetation formations in this urban forest. The general habitats of the studied amphibian were mainly in urban forests where dense forests, water bodies, and riparian vegetation were available.

**Figure 1.**
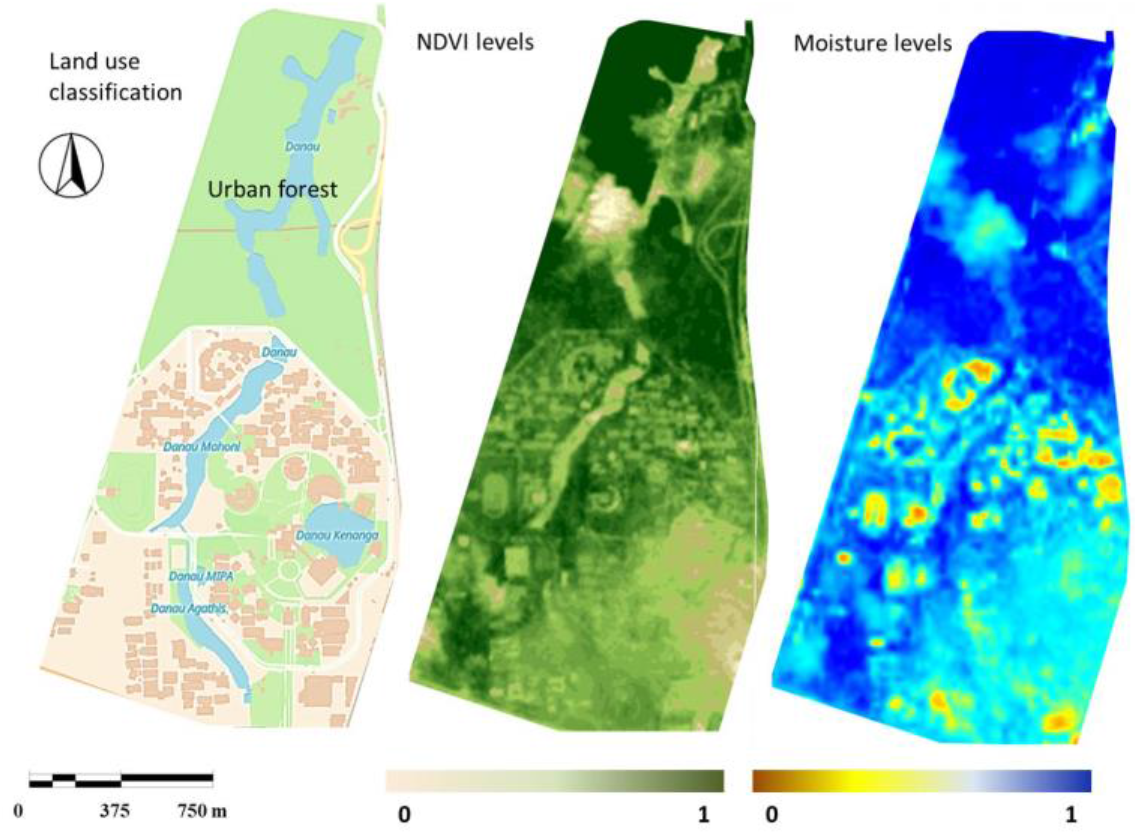
Study area (bottom left corner lon: 106.818798 E, lat: −6.373085 S and top right corner lon: 106.832929 E, lat: **-**6.349306 S), land use (developed areas, vegetation), NDVI (barren surface: 0, vegetation: 1), and moisture levels in soils and vegetation (dry: 0, wet: 1) in urban forests of Universitas Indonesia Campus in West Java.

### Occupancy (Ψ) modeling and Akaike habitat model selection

Occupancy (Ψ) modeling and Akaike habitat model selection methods were following method by Hellman (2013). Coleman et al. (2014), and Starbuck et al. (2015). In herpetology study (Mazerolle 2006), Akaike (AIC) is remarkably superior in model selection including variable selection than hypothesis-based approaches. AIC is simple to compute and easy to understand, and more importantly, for a given data set, it provides a measure of the strength of evidence for each model that represents a plausible biological hypothesis relative to the entire set of models considered. A feature of occupancy modeling is the ability to account for imperfect detection with the incorporation of detection covariates. The occupancy analysis was performed upon comparisons of events with presences of amphibian species and total sampling events. The occupancy variables were denoted as occupancy (Ψ) and occupancy analyses were performed to compare amphibian occupancy in study area as functions of hydrosphere covariates.

To accomplish this, there is a two-stage approach. First, a series of candidate detection models were created to explain varying detection probabilities for 4 amphibian species including *Bufo melanosticus, Hylarana nicobariensis, Fejervarya limnocharis*, and *Polypedates leucomystax*. The models ψ(.),p(.), which contain an occupancy ψ(.) component and a detection, or p(.) component were developed following work by MacKenzie et al. (2002). The models were tested using multi-model inference developed by Hines (2006). The top model was selected to populate the detection, or p(.), portion of the occupancy models. Then, a model set was proposed to explain occupancy of a species. The model set contained both a null and a global model. Vegetation cover tests the impact of terrestrial vegetation cover surrounding water body on amphibian occupancy. Water body tests the impact of wetland, vernal pond, and lake presences on occupancy. Riparian vegetation tests the influence of proximity and presence of aquatic and riparian vegetation cover surrounding water body on occupancy.

Occupancy model as functions of vegetation cover, water body, and riparian vegetation was developed using AIC. The AIC was developed using the linear regression. The measured parameters included in AIC are ΔAIC and AIC weight. To build the model, 3 explanatory covariates including vegetation cover, water body, and riparian vegetation and combinations of those covariates were included in the analysis to develop the model.

## Data analysis

All of the calculations and data analysis were performed using Principal Component Analysis (PCA). This analysis was employed to demonstrate which hydrosphere covariates and vegetation covers as a most covariate significantly affected the amphibian occupancy.

## RESULTS AND DISCUSSION

Based on the conducted PCA analysis (Figure 3) and correlation matrix (Figure 2) between amphibian occupancy with hydrosphere covariates and vegetation covers in urban forests of Universitas Indonesia Campus in West Java, certain correlations were observed that can be described as significant related to the formation of hydrosphere. In general, amphibian occupancy was correlated strongly with water body (r = 0.8) and riparian vegetation (r = 0.99), and show less responses on vegetation covers (Figure 4). It is clear that hydrosphere covariates are factor which strongly affects the occupancy of amphibian excepts for *B. melanosticus*. Vegetation covers were strongly influencing the *P. leucomystax*. While *H. nicobariensis* and *F. limnocharis* were influenced by the presences *of* water body and riparian vegetation. *B. melanosticus* was the only amphibian that was not correlated both with hydrosphere covariates and vegetation covers.

**Figure 2.**
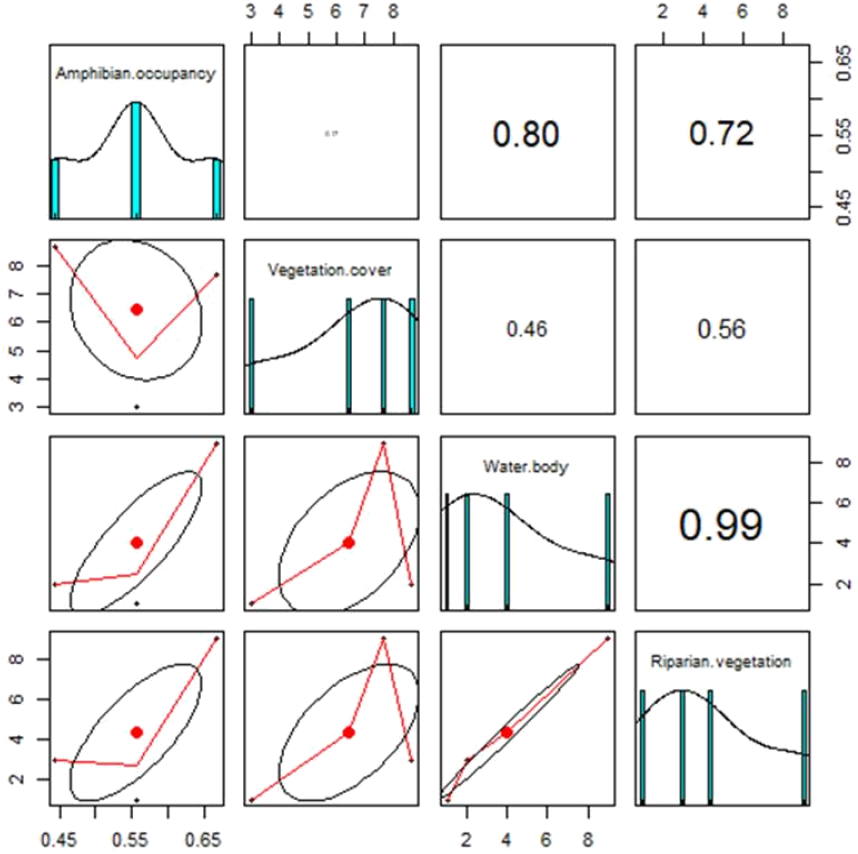
Correlation matrix of amphibian occupancy with hydrosphere covariates and vegetation covers in urban forests of Universitas Indonesia Campus in West Java.

**Figure 3.**
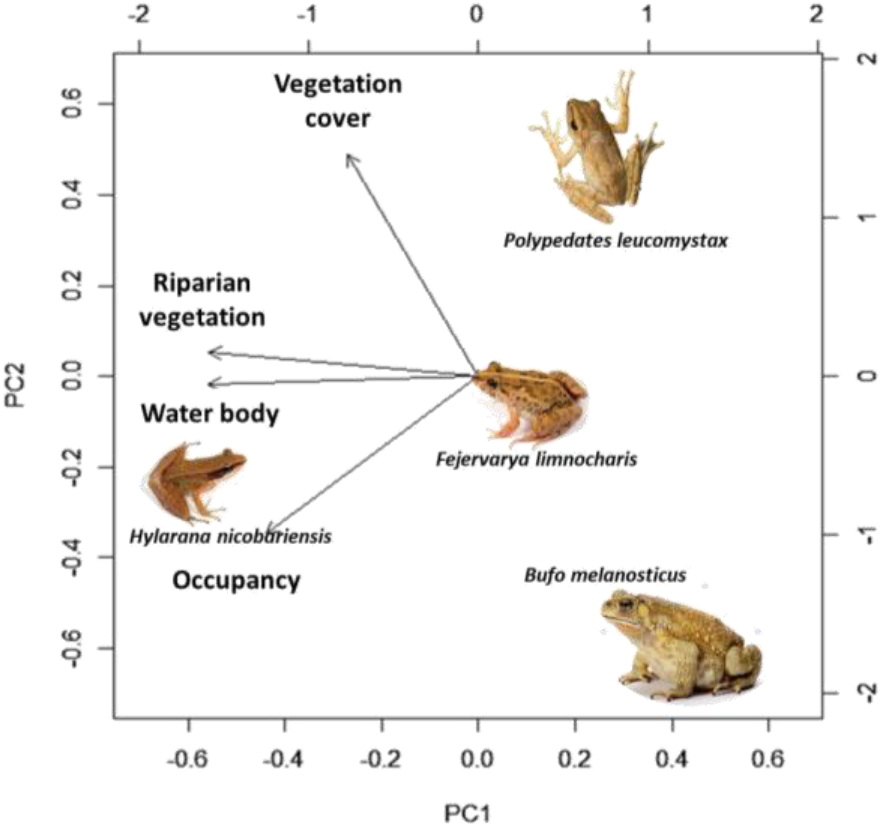
PCA of amphibian occupancy for *Bufo melanosticus, Hylarana nicobariensis, Fejervarya limnocharis*, and *Polypedates leucomystax* species with hydrosphere covariates and vegetation covers in urban forests of Universitas Indonesia Campus in West Java

**Figure 4.**
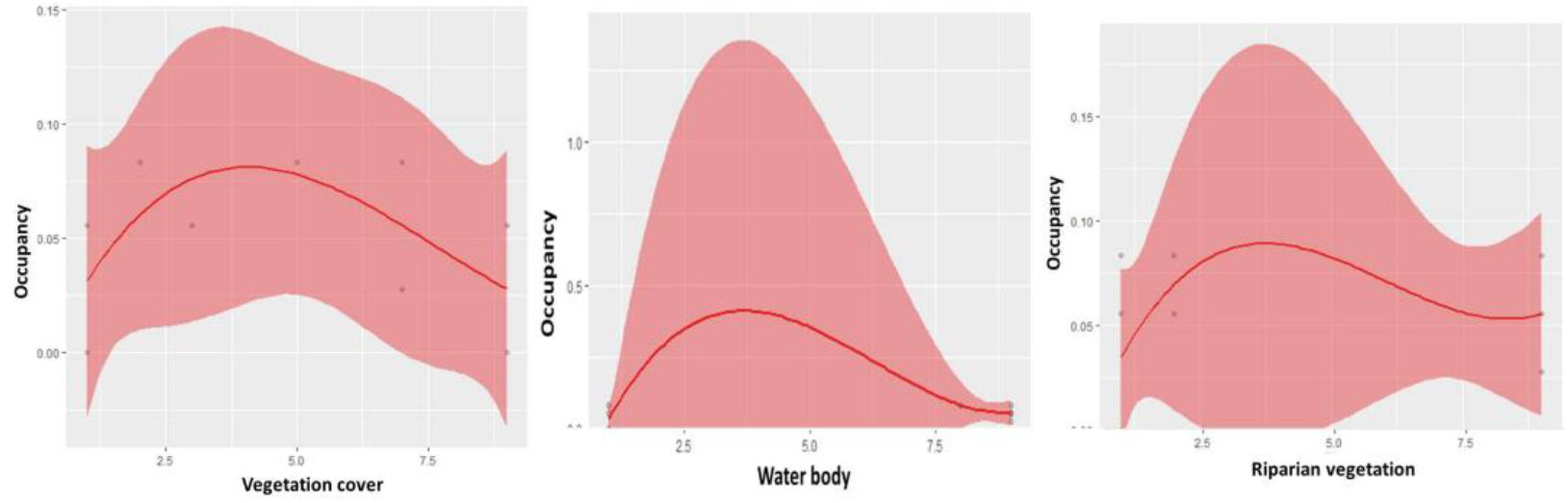
Response curves (with 95%CI in shaded areas) of amphibian occupancy (y axis) as functions of hydrosphere covariates (water body, riparian vegetation) and vegetation covers (x axis) in urban forests of Universitas Indonesia Campus in West Java

Seven occupancy models tested for the 4 amphibian species in urban forests of Universitas Indonesia Campus in West Java can be seen in Table 1. For *Hylarana nicobariensis*, hydrosphere covariates and vegetation covers have provided similar effects on this species occupancy. It means that those microhabitats all are important. *H. nicobariensis* is known as amphibian that can inhabit wide variety of microhabitat as long water body and riparian vegetation are available (Putra et al. 2012, Kurniati & Hamidi 2016). For *Polypedates leucomystax*, riparian vegetation is more important to determine habitat occupancy of this species followed by the water body. P. *leucomystax* has Ψ values of −14.18 for riparian vegetation and −12.59 for water body microhabitat. *P. leucomystax* is an arboreal frog that requires both water body combined by the presence of wetland vegetation (Muslim 2017). This vegetation is providing substrate for frog to perch. Similar to *P. leucomystax, Fejervarya limnocharis* also has high occupancy for riparian vegetation and water body with Ψ values of −12.59 and −11.64. Water body and riparian vegetation are important covariates for amphibian considering water bodies are required to reproduce, and a riparian vegetation are also needed where some amphibian species live outside their breeding period (Mazgajska & Mazgajski 2020).

**Table 1.** Seven occupancy models tested for the 4 amphibian species in urban forests of Universitas Indonesia Campus in West Java (asterisk sign show the best models).

| Amphibian species occupancy ($\Psi$) + covariate models | AIC | ΔAIC | AIC weight |
| --- | --- | --- | --- |
| *Hylarana nicobariensis* |  |  |  |
| $\Psi$(vegetation cover) | -6.54* | 0 | 0.33 |
| $\Psi$(water body) | -6.54* | 0 | 0.33 |
| $\Psi$(riparian vegetation) | -6.54* | 0 | 0.33 |
| $\Psi$(vegetation cover+water body) | -0.54 | 0 | 0.30 |
| $\Psi$(vegetation cover+riparian vegetation) | -0.54 | 0 | 0.30 |
| $\Psi$(riparian vegetation+water body) | -0.54 | 0 | 0.30 |
| $\Psi$(vegetation cover+water body+riparian vegetation) | 2 | 0 | 0.11 |
| *Polypedates leucomystax* |  |  |  |
| $\Psi$(vegetation cover) | -10.34 | 3.84 | 0.09 |
| $\Psi$(water body) | -12.59* | 1.59 | 0.28 |
| $\Psi$(riparian vegetation) | -14.18** | 0.00 | 0.63 |
| $\Psi$(vegetation cover+water body) | -8.18 | 0 | 0.3 |
| $\Psi$(vegetation cover+riparian vegetation) | -8.18 | 0 | 0.3 |
| $\Psi$(riparian vegetation+water body) | -8.18 | 0 | 0.3 |
| $\Psi$(vegetation cover+water body+riparian vegetation) | -6.18 | 2 | 0.11 |
| *Fejervarya limnocharis* |  |  |  |
| $\Psi$(vegetation cover) | -10.36 | 2.23 | 0.17 |
| $\Psi$(water body) | -11.64* | 0.95 | 0.32 |
| $\Psi$(riparian vegetation) | -12.59** | 0.00 | 0.51 |
| $\Psi$(vegetation cover+water body) | -5.64 | 2.36 | 0.16 |
| $\Psi$(vegetation cover+riparian vegetation) | -4.36 | 3.63 | 0.09 |
| $\Psi$(riparian vegetation+water body) | -8.00 | 0.00 | 0.53 |
| $\Psi$(vegetation cover+water body+riparian vegetation) | -6.17 | 1.82 | 0.21 |
| *Bufo melanosticus* |  |  |  |
| $\Psi$(vegetation cover) | -14.18* | 0.00 | 0.88 |
| $\Psi$(water body) | -8.79 | 5.39 | 0.06 |
| $\Psi$(riparian vegetation) | -8.79 | 5.39 | 0.06 |
| $\Psi$(vegetation cover+water body) | -8.18 | 0.00 | 0.41 |
| $\Psi$(vegetation cover+riparian vegetation) | -8.18 | 0.00 | 0.41 |
| $\Psi$(riparian vegetation+water body) | -2.79 | 5.39 | 0.03 |
| $\Psi$(vegetation cover+water body+riparian vegetation) | -6.18 | 2.00 | 0.15 |

*B. melanosticus* was the only amphibian species that has low occupancy on hydrosphere covariates with Ψ value of −8.79 and high occupancy for terrestrial vegetation cover (Ψ value = −14.18). This finding is in line with the majority of the other studied urban landscapes. *Bufo* is known as genus to be adaptable to heterogeneous habitats, including urban areas (Pavignano et al. 1990, Berger 2008, Mollov 2011). Budzik et al. (2013) noticed that *Bufo* was the only amphibian which increased, while the others declined in urban landscapes. These findings may suggest the high resistance of *Bufo* to the negative impact of urbanization including loss and fragmentation of habitat and pollution.

In urban forest, *Bufo* was more common in areas near settlements while other amphibian species were observed in the isolated hydrosphere far from settlements. Hamer & Parris (2011) found that amphibian species richness decreased at hydrosphere surrounded by high densities of human residents and amphibian richness increased substantially at hydrosphere surrounded by a high proportion of green open space and riparian vegetation. Urbanization had strong negative effects on amphibian species that were associated with well vegetated water bodies.

This study is the first in South East Asia regions that provides empirical evidences of amphibian occupancy related to hydrosphere covariates mainly in urban settings. The findings in this study were in agreement with other studies. The permanency of water bodies, their occurrence in the vicinity of river valleys, and a high ratio of riparian vegetation around water bodies are positively correlated and have a significant influence on amphibian occupancy within the hydrosphere of urban landscape. Thus, these identified factors should be considered in the course of sustainable urban planning in order to avoid potential conflicts between nature conservation, hydrosphere sustainability, and urban development.

